# A Cortico-Cerebellar Network Model for Refining Preparatory Activity in Motor Control through Sensorimotor Learning

**DOI:** 10.64898/2026.08.10.743900

**Authors:** Serhat Çağdaş, Neslihan Serap Şengör

## Abstract

This paper introduces a sensorimotor learning framework for a corticocerebellar network, grounded in the perspective of population dynamics. Using an optimal control theory approach, the cerebellum model enhances preparatory activity through premotor input, allowing the motor cortex to reach the desired initial conditions for movement more efficiently. Unlike traditional motor learning approaches that focus on acquiring new skills, this paradigm emphasizes automatization of already executable behaviors through repetition driven by intrinsic motivation. The proposed model is evaluated using a center-out reaching task, demonstrating that the role of the cerebellum is to shorten the preparatory period required for the successful execution of the movement. These findings suggest that corticocerebellar interactions play a crucial role in optimizing motor preparation, offering insight into the neural mechanisms underlying movement efficiency.

## I. Introduction

The encoding of movement characteristics, e.g. velocity and force, by neurons of the motor cortex, the primary controller of the descending motor system, has been a long-standing topic of research [1], [2]. However, in recent years, the perspective of population dynamics has gained momentum against the single-neuron doctrine, which is based on the idea that nerve cells are discrete information processors [3].The dynamical system perspective claims that population encoding is essential rather than a single-neuron representation, although there can be a correlation between individual neural activity and movement kinematics [4].

Movement through a population dynamics perspective can be summarized in three key points [5], [6]:

- The motor cortical population’s behavior constitutes a rotational dynamics, with a dimension much lower than the number of neurons.
- The movement generated heavily depends on preparatory activity that drives the population toward an initial condition associated with the movement.
- Activities related to preparation and movement time exist in orthogonal subspaces.

The new ideas on motor control by population coding require also a re-assessment of the motor learning paradigms and how subcortical areas such as the basal ganglia and cerebellum contribute to motor learning [7].

The basal ganglia are believed to contribute to action selection and sequential learning, whereas the cerebellum, regarded as a supervised learning center of the nervous system, plays a more significant role in sensorimotor adaptation [8], [9]. However, how action selection and refinement of motor behavior is achieved in the perspective of population dynamics is another question. As an example, [10] proposed that the preparation and execution of each motif in a sequence are selected by the basal ganglia and the motifs are chained using the basal ganglia as a switch.

In this study, we focus on sensorimotor learning and the contribution of the cerebellum to motor control by enhancing the efficiency of preparatory activity. In experimental studies, it is proposed that the cerebellum is involved in motor planning, adaptation, and timing by affecting preparatory activity [11]– [13]. In another study on patients with unilateral cerebellar stroke, it is suggested that lesions in the cerebellum may induce deficits in movement preparation [14].

In a previous model, the duration of preparatory activity required for proper reaching behavior was reduced by optimizing feedback input using a linear quadratic regulator algorithm [15]. Here, we propose a bio-inspired solution involving corticocerebellar sensorimotor learning. In a center-out reaching task, the two joint-arm model controlled with a corticocerebellar network is supposed to repeat the movement behaviors that it has already successfully achieved with long preparatory activity durations. As the system continues learning through cerebellar plasticity, it converges the cortical conditions necessary for movement with shorter preparatory activity and reaches the targets successfully after short preparatory activities.

## II. Related work

After Marr’s associative motor memory model with the cerebellar cortex, different computational frameworks have been proposed for sensorimotor learning in the cerebellum [16]. In these motor learning frameworks, the cerebellum is typically characterized as a forward or inverse internal model [17], [18]. Cerebellar mechanisms effective in optokinetic eye movement response [19], vestibulo-ocular reflex [20], and saccadic eye movement adaptation [21] are widely investigated using the idea of predictions from these internal models.

In the majority of these studies, the cerebellum is modeled as a controller of the object or descending pathway exclusively, rather than contributing to the motor control of the cerebral cortex. In another computational approach, the cerebellum cooperates with the basal ganglia and the motor cortex to control a two-joint arm [26]. However, in this case, the cerebellar signal is used to adjust movement, rather than modulating motor cortical activity. On the other hand, in the studies that recruit the cerebellum as a feedback decoupling machine for the motor cortex, the projection weights of the lower-dimensional cerebellar network to the motor cortex must also be optimized as part of learning [27], [28].

It is also proposed that the cerebellum is involved in subsecond temporal perception and response. A computational model is used for the delayed eye blink conditioning response task [22]. The delay line [23] and spectral timing models [24] are used to mimic the temporal process in the cerebellum. Dean et al. [25] state that the cerebellar microcircuit processes information as an adaptive filter, facilitated by recurrent connections in the mossy and parallel fibers.

## III.Proposed Cortico-Cerebellar Framework

In this study, the cerebellum is considered as an inverse internal model. It gets sensory input and generates a response taking advantage of the temporal basis set generated by the granular layer. The cerebellar network modulates premotor activity to help the primary motor cortex converge to the desired initial condition 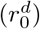 faster during the preparatory activity period (Fig. 1).

**Fig. 1.**
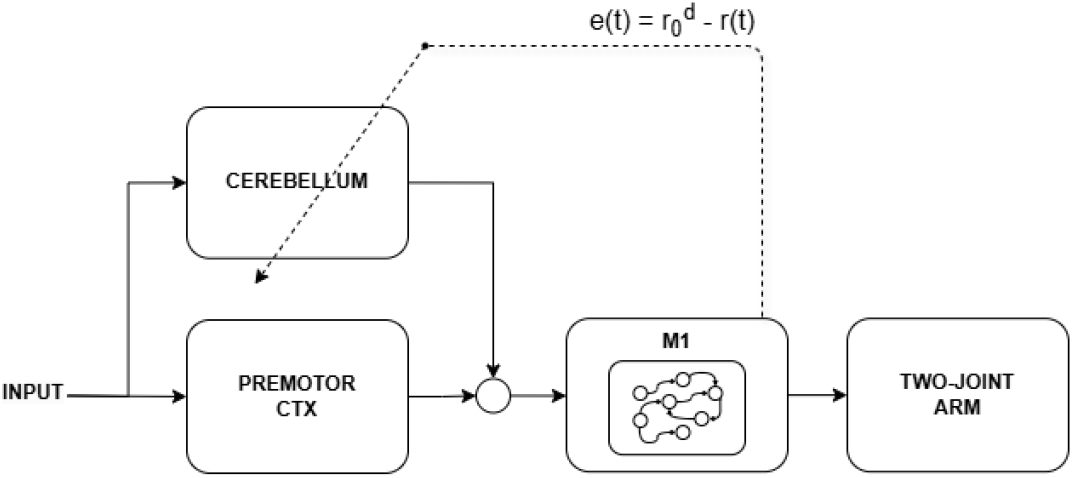
Corticocerebellar network diagram.

The primary motor cortex is modeled as a recurrent dynamical system consisting of *N* = 100 rate-based neural units:

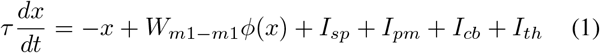

where *x* is the internal activation of the neural units. *τ* is the time constant of the units and is set as 100 ms for the whole primary motor cortex. The frequency rate of each unit is a non-linear function of internal activation and is represented by *r*_*i*_ = *ϕ*(*x*_*i*_). In this study, it is used as *ϕ*(*x*_*i*_) = *ln* 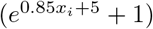. Related to the non-linear rate function, *I*_*sp*_ = − *W*_*m*1*−m*1_*ϕ*(0) generates a spontaneous cortical activity around 5*Hz*.

The recurrent structure of M1 is achieved with the internal connection matrix *W* which is a linear combination of the symmetric and skew-symmetric components of a random matrix *M* where *M*_*ij*_ *~ U*(−0.5, 0.5) [30]:

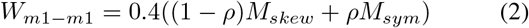

Therefore, M1 can show transient quasi-rythmic activity in response to simple input. In (2), *M*_*skew*_ = (*M* − *M* ^*T*^)*/*2 and *M*_*sym*_ = (*M* + *M* ^*T*^)*/*2 are skew-symmetric and symmetric matrices, respectively. The constant *ρ* is determined as 0.25. Therefore, the imaginary part of the eigenvalues is tuned in relation to the proportion of the skew-symmetric component, and the RNN model used as M1 can show rotational transient trajectories as a dynamical system.

Finally, *I*_*pm*_, *I*_*cb*_ and *I*_*th*_ are the input provided by the pre-motor cortex, the cerebellum, and the thalamus, respectively, in (1). Thalamic input *I*_*th*_ = *g*_*bg*_*r*(*t*) is used to amplify cortical activity. The input is an abstract representation of the cortico-basal ganglia-thalamus feedback where *g*_*bg*_ is the gate variable representing the basal ganglia and it gets 5 or 0 values in this study. On the other hand, the premotor input for each M1 neural unit is simply simulated with a linear transformation of sensory input:

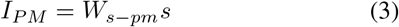

where The vector *s ∈* ℝ^5^ represents the data obtained from the sensor. A topographically organized connection matrix is chosen for *W*_*s*_→*pm* as in [31]:

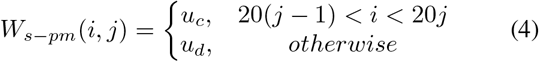

where *u*_*c*_ *∈ U* [0.5, 1.2] and *u*_*d*_ *∈ U* [0, 0.2] are uniformly distributed random values.

On the other hand, a feedforward structure is used to model the entire cerebellum (Fig. 2). The sensory input of the network is provided by the mossy fibers (*MF* = *s ∈* ℝ^5^). The deep cerebellar nucleus (*DCN ∈* ℝ^*N*^), which is the output layer of the cerebellum, is composed of *N* units. It is excited by the mossy fibers and inhibited by Purkinje cells (*PC ∈* ℝ^*N*^):

**Fig. 2.**
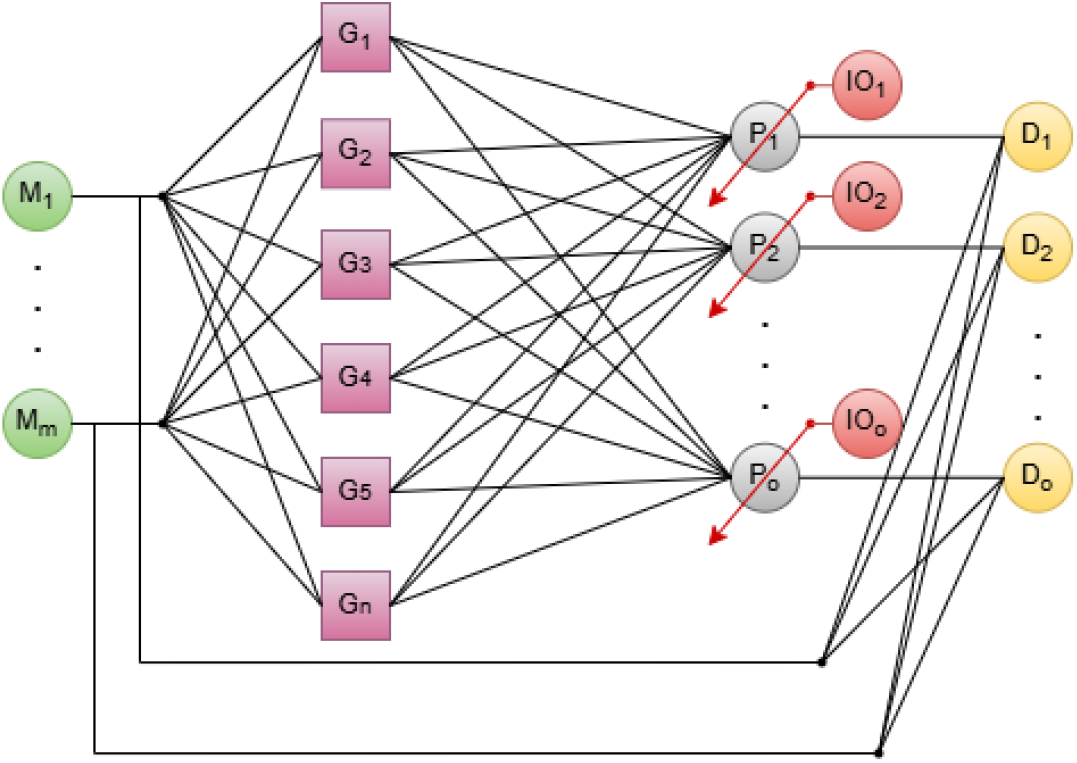
Feedforward cerebellum structure consisting of mossy fiber (M), Granular (G), Purkinje (P) and deep cerebellar nucles(D) layers. Inferor olive (IO) sends instructive signal to P units. M-G, G-P and M-D connections are all-to-all, whereas P-D and P-D are connected in a one-to-one manner.

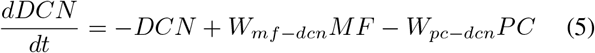

where *W*_*mf−dcn*_(*i, j*) = 2 *∈* ℝ^5*XN*^ and *W*_*pc−dcn*_ is a unity matrix of dimension *I ∈* ℝ^*NXN*^. The granular layer stimulates the PC units, which in turn indirectly suppress the DCN:

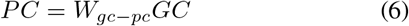

The granular layer is represented with a temporal basis set consisting of heterogeneous Gaussian kernels as in [32]:

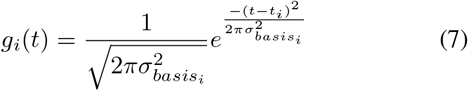

The kernel that belongs to granular unit *GC*_*i*_ accross time *g*_*i*_(*t*) reaches its peak value at time *t*_*i*_ after the start of preparatory activity (Fig. 3). The value of each unit is determined as *t*_*i*_ = 5 + (*t*_*max*_ − 5)(*i/N*_*gc*_) where *t*_*max*_ = 1300 ms and the number of granular units is *N*_*gc*_ = 2500. The width of the kernels are also determined depending on index of the units as 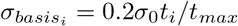 The parameter *σ*_0_ is set as 500 ms during simulations.

**Fig. 3.**
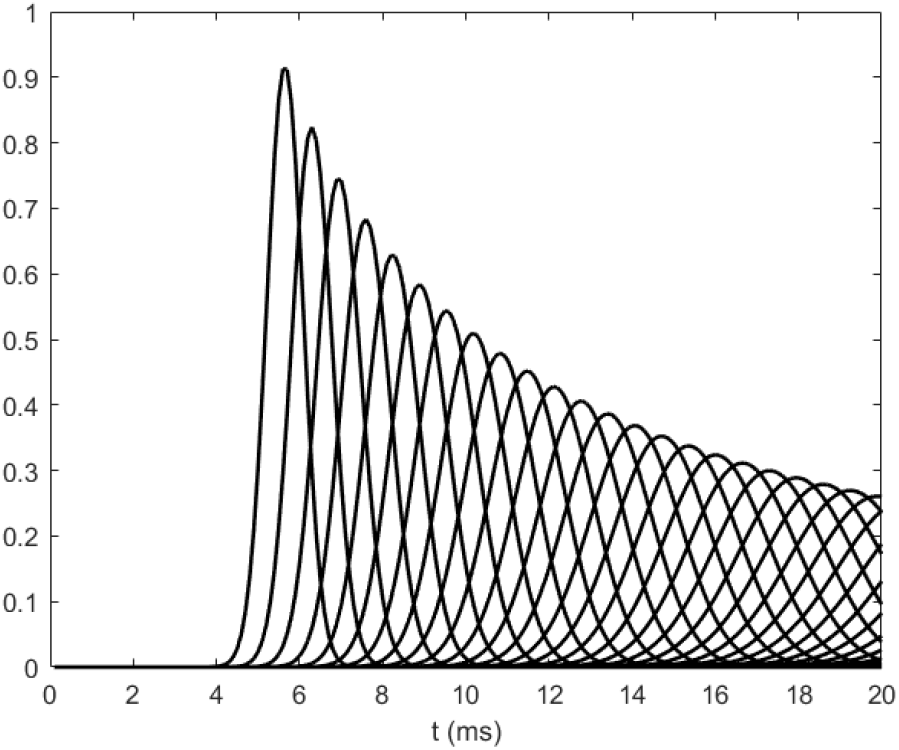
Heterogeneous temporal basis kernels of granular units.

The temporal amplitude of the activity of units of the granular layer depends on the kernel *g*_*i*_(*t*) and the sensory input conveyed through MFs:

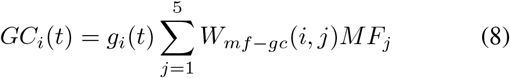

The connections between the granular layer and the Purkinje layer are plastic. *W*_*gc−pc*_ connections are updated due to instructive signals coming from the inferior olive neurons (IO) and the activities of granular neurons. IO units generate binary error signals depending on the bias of the preparatory activity of M1 units from the desired initial conditions.

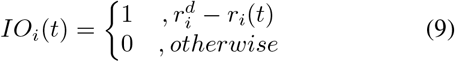

In this basic plasticity rule, granule cell activities lead to long-term depression (LTD) in the presence of IO activity. In contrast, long-term potention occurs in the absence of IO input [20], [22]:

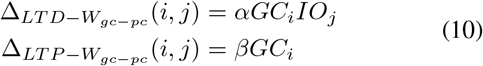

*α* = 0.002 parameter is the plasticity rate for LTD and *β* = 0.001 parameter is the plasticity parameter for LTP. In addition to LTP and LTD updates, a recovery plasticity component is included that helps weights return slowly to their initial values:

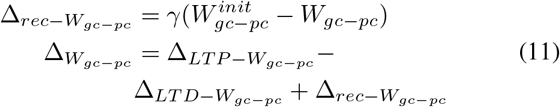

where the parameter *γ* = 10^*−*6^ is the convergence rate of the recovery component. The weights of all *GC*-*PC* synapses are initialized to 1 and it is ensured that they remained in the [0, 10] interval throughout training.

## IV. EXPERIMENTAL PARADIGM

The network is tested on a center-out reaching task. The two-joint arm model in [29] is used as the controlled object during simulations. At the beginning of each trial, the two-joint arm is positioned at the idle joint angle (*θ*_*joint*1_ = 10, *θ*_*joint*2_ = 143.54). There are five targets that are placed equi-distant (*d* = 0.2*m*) from the idle point and target *i* is in the direction of the angle (0.4*i* − 0.7)*π*. One of the targets are shown during the preparatory activity. The sensory input related to the current target is set to *s*(*i*) = 5.

At the end of the preparation period (*t* = *T*_*prep*_), a start cue is shown and thalamic input is switched on (*I*_*th*_ = 5*r*(*T*_*prep*_)) for a duration of 50*ms*. Following the thalamic input, reaching behavior is executed in the consequent 1000*ms* duration (Fig. 4). The torque data for each joint *m*(*t*) *∈* ℝ^2^ motor execution are generated by the internal transient pattern of the recurrent M1 network:

**Fig. 4.**
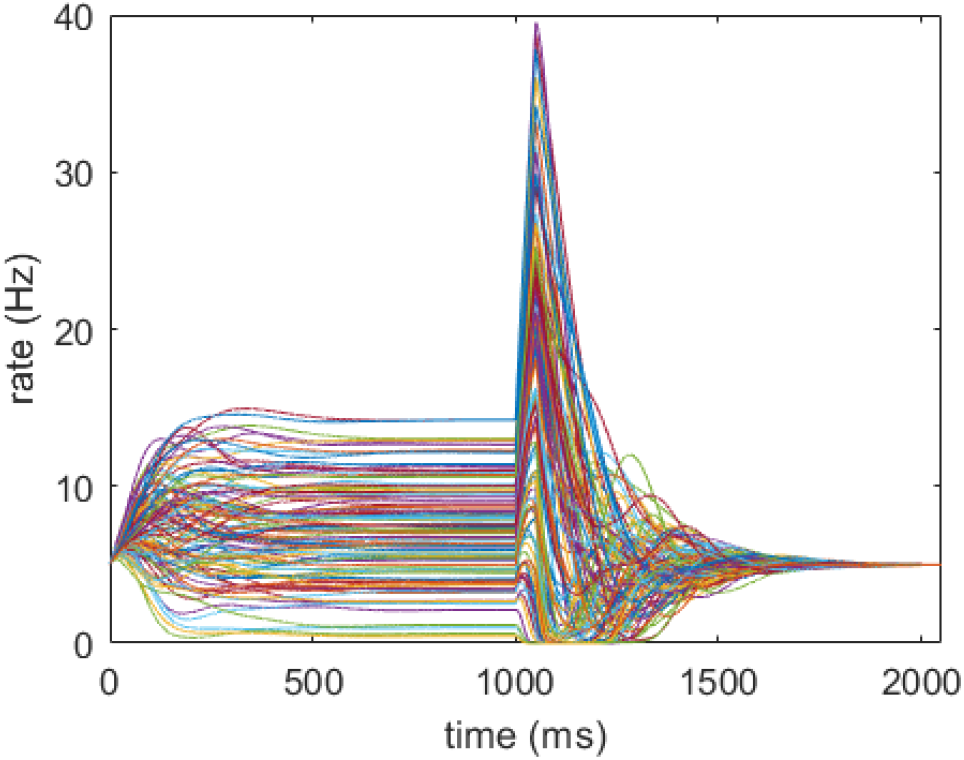
One trial cortical activity with a preparatory time of 1000 ms. Phasic thalamus input lasts for 50 ms and remaining 1000 ms run is left as execution time.

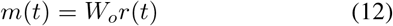

where *W*_*o*_ *∈* ℝ^*Nx*2^ is the read-out matrix of the M1 population.

### A. Read-out Matrix Optimization

For optimization of the read-out matrix, the Tikhonov regularization method is used [30], [33].

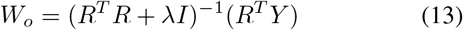

*R ∈* ℝ^5*TxN*^ is the concatenated matrix composed of cortical activities under five different sensory inputs and reach trials. The preparatory activity duration is determined as 1000*ms*. Therefore, M1 has enough time to reach its equilibrium state as it is influenced by the premotor area, which is driven by sensory input. In addition, optimization has been performed for the case in which the cerebellum is ablated from the network. For this reason, *I*_*cb*_ *∈* ℝ^*N*^ denotes input with all components set to 5 Hz during the prepatory activity in optimization process, representing the average activity of the DCN units.

Another matrix, *Y ∈* ℝ^5*Tx*2^ in (14) is the concatenated time series of the desired torque data to reach five targets. *T* is 20500 for both matrices considering the 1000*ms* preparatory, 50 ms thalamic input and 1000 ms movement execution period with timestep *dt* = 0.1 ms. Reaching activity is only allowed at execution period. So, *m*(*t*) is 0 when 0 < *t* < 1050 ms. Linear arm trajectories are aimed for reaching [34] and, the desired torque data for each target are obtained with a proportional derivative feedback controller using the parameters in [35].

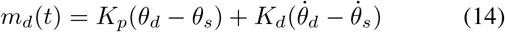

*θ*_*d*_ and *θ*_*s*_ are the desired and spontaneous angular vectors. The controller parameters are *K*_*p*_ = 100 and *K*_*d*_ = 1.

## V. RESULTS OF CENTER-OUT REACHING TASK

After regression of the read-out matrix without the cerebellum case, the equilibrium states of M1 in the preparation period configuration were saved as desired initial conditions (*r*^*d*^) for different directions. These initial conditions were used to generate the IO instructive signals as in (9). The cerebellum was trained with 500 trials with random target inputs and a preparatory activity duration of 1000 ms.

The mean square error of the difference between the desired initial condition and the spontaneous activity of M1 is calculated as the activity bias throughout the preparatory activity period. Fig. 5 shows the first 500 ms of preparatory activity of the M1, PC, and DCN populations and the activity bias value of M1 for one trial. The default activity bias is observed in Fig. 5a when the cerebellum is ablated. After including the cerebellum, the M1 activity bias values increase slightly, since the PC and DCN populations add perturbation to the preparatory activity of M1 (Fig. 5b). Contrary to this negative effect in the first trial, the cerebellum refines preparatory activity and causes M1 to converge the desired initial condition faster after the training procedure as expected (Fig. 5c).

**Fig. 5.**
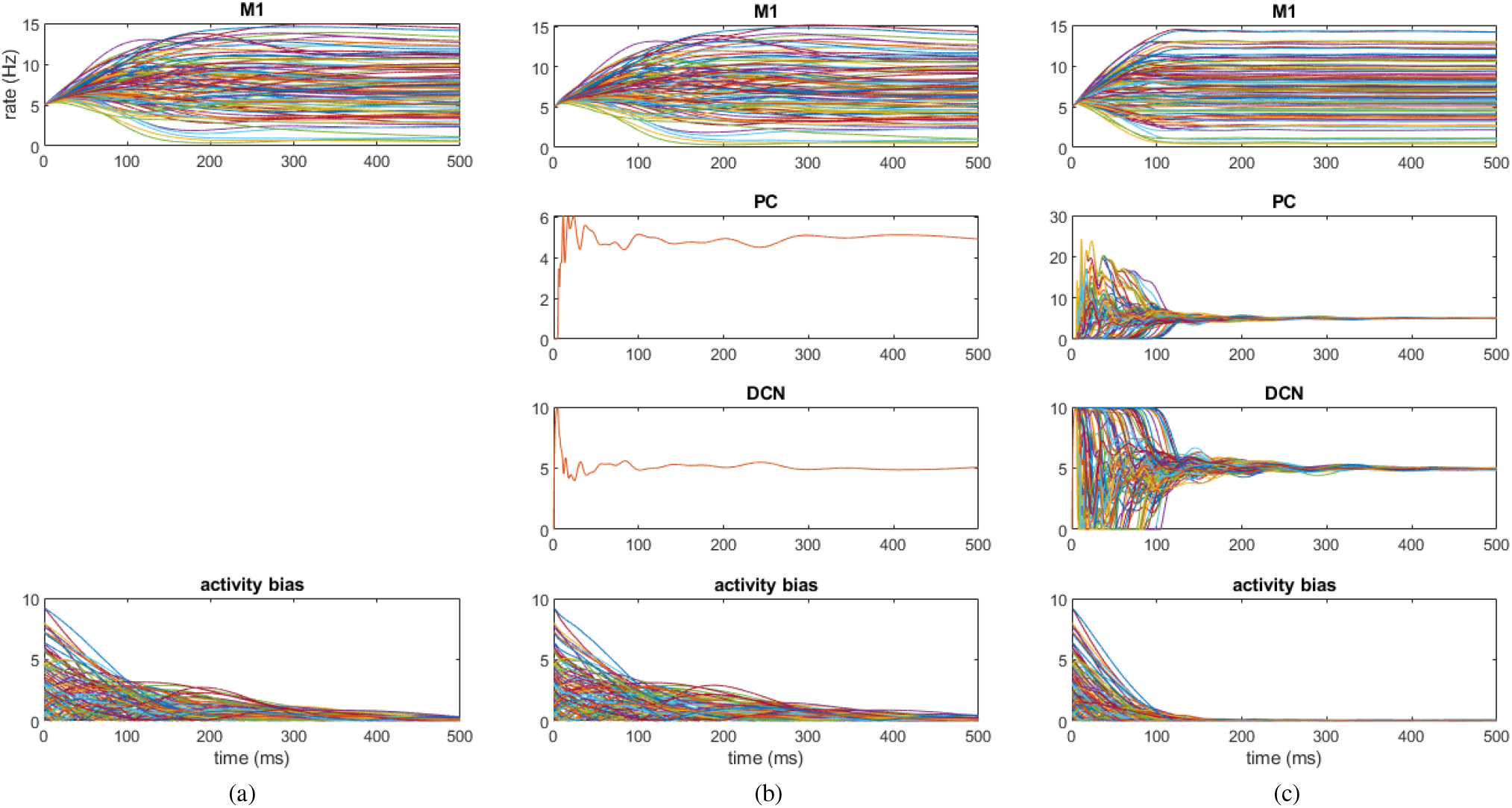
M1, PC and DCN activities for first 500 ms of the preparatory activity period in (a) a trial for the ablated cerebellum network, (b) first trial and (c) 500th trial of cortico-cerebellar network. Bias values from the desired initial conditions through time are given at the bottom.

Reaching trajectories for five targets are also observed with the same network configurations. To prevent drift on the trajectory *m*(*t*) is set artificially to 0 during preparatory activity as in [34]. Fig. 6a shows the reaching trajectories with a preparatory duration of 1000 ms when the cerebellum is absent. The data generated by cortical activity in those trials are also used for read-out matrix optimization. When the reaching task is repeated with 300 ms preparatory activity, reaching the targets is failed (Fig. 6b). This is mainly because the initial condition for the corresponding reaching behavior cannot be converged by M1 in such a short time as observed in Fig. 5a. A similar failure is encountered in the network with an untrained cerebellum and preparatory activity of 300 ms (Fig. 6c). However, once the training is finished a 300 ms preparatory activity is sufficient for successful reaching behaviors anymore.

**Fig. 6.**
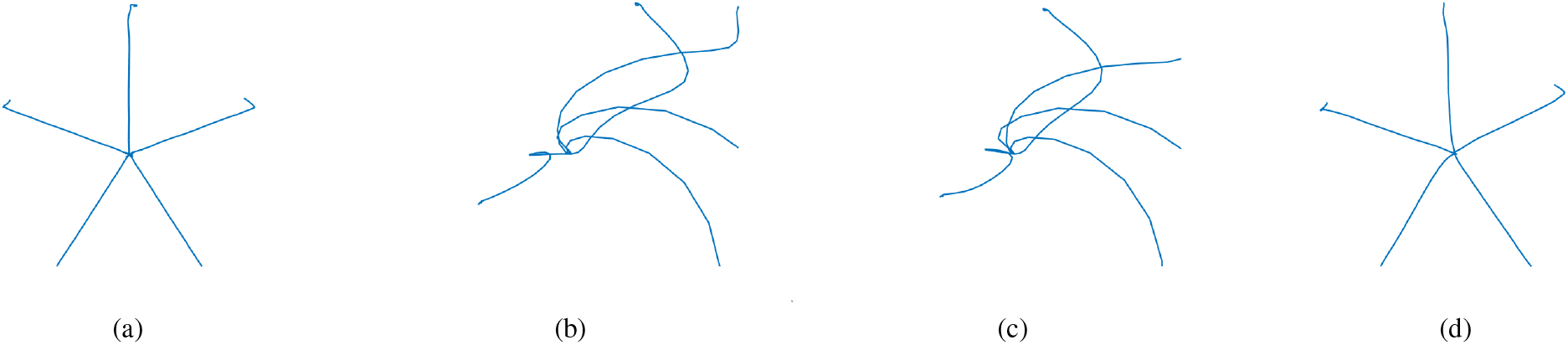
Reaching trajectories of the ablated cerebellum network with a preparatory period of (a) 1000 ms and (b) 300 ms. (c) and (d) are the trajectories of corticocerebellar network with a period of 300 ms in the first and 500th trial, respectively.

Finally, the task is repeated for a set of preparatory durations and prospective motor error values (*ϵ*_*p*_) are calculated for further analysis. The ratio of the energy of the difference between the generated and desired torque data to the energy of the torque data has been used for error analysis.

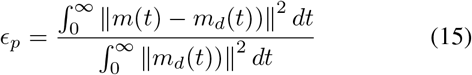

The results in Fig. 7 are consistent with the previous tests. In the ablated cerebellum case, the reaching behavior can be executed plausibly only after a preparation time of 400 ms. When the cerebellum is added, a small bias is observed from the default network results because of the DCN perturbation. After training, reasonable results are encountered with a preparation time of more than 150 ms.

**Fig. 7.**
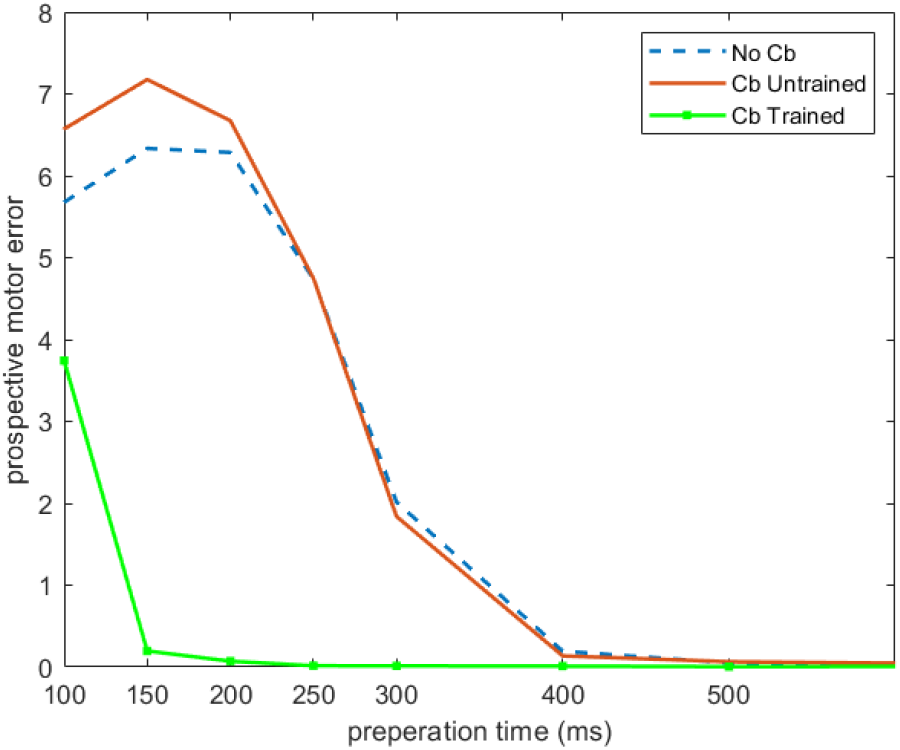
Prospective motor error analysis for the ablated cerebellum (dashed line), untrained (solid line) and trained (dotted line) cortocerebellar networks.

## VI. DISCUSSION

In this work, we present a sensorimotor learning paradigm for a corticocerebellar network, aligned with the population dynamics perspective. Within an optimal control theory framework, the cerebellum learns to refine preparatory activity, enabling the motor cortex to achieve the desired initial conditions for movement in a shorter duration.

There are two main parts of the network: a recurrent neural network as the primary motor cortex and a feedforward cerebellar structure. The RNN as a dynamical system receives the input related to sensory data from an abstract premotor area and the cerebellum in the preparatory activity period. After converging the desired initial condition, it transiently generates a quasi-rythmic pattern related to movement by its internal dynamics in the execution time.

In the later stages, the cerebellum receives the cortical activity bias for the desired initial condition via the inferior olive (IO) as an instructive signal. Parallel fiber synapses are then updated to minimize this error. At this point, the basis kernels in the granular layer facilitate temporal learning.

It should be noted that the sensorimotor learning process refines behavior by reducing the duration required for preparatory activity rather than acquiring a new motor skill. This type of learning is more related to the automatization of behaviors that can already be performed. Automatization is achieved through repetition driven by intrinsic motivation. Typing on a keyboard is a simple example of this type of automatization. On the other hand, the process may also contribute to sequence learning, as shorter preparatory motifs are required between the execution motifs in this way [10].

In the study, number of granular layer units is limited to 2500 considering the simulation time. However, it was observed that the prospective motor error is tightly related to the number of granular cells. Although the current configuration results in successful reaching trials with preapratory period greater than 150 ms, the learning success may decrease in case of increasing the number of targets. Then, using a larger granular layer may solve the problem.

Secondly, temporal basis kernels that represent granular units may affect the performance of the network. Here, a temporal basis set was selected in a way that the DCN mean activity remains on a steady state during preparation for simplicity. However, Narain et al. [32] investigated different parameters of the Gaussian kernel for a timing task and concluded that these parameters affect the performance of the network.

Furthermore, the only plastic synapses of the network belong to the GC-PC connections in this study. There are known to be other connections that are plastic and contribute to learning [37]. In [20], plasticity of the MF nucleus connection is suggested to consolidate the VOR adaptation. This plasticity may provide a short latency response for the early preparatory activity in this work as proposed in [22].

In the network, the only dynamical part of the cerebellum was the deep cerebellar nucleus. Since it has a very small time constant, the structure contributes to the cerebral cortex with almost no delay. The success of refinement under delay conditions remains an open question for the network and needs to be further analyzed. Another direction of the work may also be enhancing the performance of the network in a readaptation task in case of the change in the sensory - movement association, etc. [36].

Finally, as mentioned above, this study deals with the refinement of skills that are already in the repertoire. Therefore, read-out matrix optimization of primary motor cortex is not part of sensorimotor learning. In addition, the peripheral nervous and musculoskeletal system is outside of the scope of this study.

